# Early developmental characterization of gap-induced prepulse inhibition and habituation in the Fragile X Syndrome mouse model

**DOI:** 10.1101/2025.10.13.682100

**Authors:** Abdullah Abdullah, Xiuping Liu, Kartikeya Murari, Jun Yan, Ning Cheng

## Abstract

**Introduction:** Atypical sensory processing, particularly auditory hypersensitivity, is a common and debilitating phenotype of Fragile X Syndrome (FXS). Electrophysiology studies in *FMR1-*knockout (KO) mice previously observed hyperexcitability at the auditory cortex (AC), with enhanced neuronal firing to auditory stimuli. Prepulse inhibition (PPI), a behavioral measure of sensorimotor gating, is robustly impaired in FXS individuals. Interestingly, a related paradigm called gap-induced inhibition of the acoustic startle (GPIAS) is mediated by the AC, and a previous study observed decreased GPIAS in mature *FMR1-*KO mice. Further, habituation is also an important sensory filtering mechanism, which has been reported to be impaired in mature *FMR1-*KO mice. However, not much is known about GPIAS and habituation in *FMR1-* KO mice during early development.

**Methods:** We evaluated GPIAS in male and female *FMR1-*KO mice at post-natal days 15 (P15), 20 (P20) and 30 (P30). The paradigm consisted of a prepulse stimulus (a gap in a continuous background noise) followed by a startle stimulus, with an inter-stimulus interval of 50 or 100 ms. Habituation was assessed prior to the GPIAS trials, with a series of startle only stimulus.

**Results:** We observed a trend for genotype difference in acoustic startle response (ASR) magnitude, particularly in the female mice at P30. No significant genotype differences were noted in GPIAS or latency of ASR. Response duration was significantly increased in the male *FMR1*-KO mice compared to their WT counterparts during early development. Significant genotype differences were also observed in habituation and sensitization. In terms of development, we observed a significant increase in GPIAS with maturation. Furthermore, we also observed significant changes in the magnitude, response latency and duration of ASR with maturation. Finally, significant sex differences were observed in ASR magnitude and duration.

**Conclusion:** Our findings suggest that behavioral responses to auditory stimuli are dynamic during development and differ between females and males, an important consideration for future study design. Additionally, the *FMR1-*KO mice display habituation deficits during early development, which could be an ideal window for addressing auditory hypersensitivity in FXS.

## Introduction

Fragile X Syndrome (FXS) is the leading inherited cause of autism spectrum disorder (ASD), with an estimated prevalence of 1 in 4000 males and 1 in 8000 females [1, 2]. In general, the number of CGG trinucleotide repeats, located at the 5’ untranslated region of the Fragile X Messenger Ribonucleoprotein 1 (*FMR1*) gene, ranges between 6 to 55 repeats. An augmentation in the number of CGG trinucleotide repeats leads to a full mutation (>200), subsequently causing hypermethylation at the CpG site of the promoter region followed by the silencing of the *FMR1* gene [3, 4]. As a result, the expression of the Fragile X Messenger Ribonucleoprotein (FMRP) is disrupted leading to various FXS symptoms such as intellectual disability, anxiety, social deficits and language impairments [1, 5].

Auditory hypersensitivity or hyperacusis, presenting as decreased sound tolerance, is another common and debilitating phenotype in ASD and FXS [6-9]. Interestingly, in one study by Amir et al, 60% of the 412 children seeking care for hyperacusis had previous diagnosis of ASD [10]. This phenotype can also be observed in the *FMR1-*knockout (KO) mice, a well-established preclinical mouse model of FXS [11, 12], besides other FXS-modelling phenotypes such as hyperactivity [11] and alterations in dendritic spine shape and density [13]. Previous event-related potential (ERP) studies observed increased N1 and P2 amplitudes in FXS individuals [14, 15], although only increased N1 amplitude was displayed in the *FMR1-*KO mouse model [16]. Moreover, electrophysiology recordings at the inferior colliculus (IC) of the *FMR1-*KO mice also noted increased neuronal firing in response to tones of different frequency and amplitude [17]. However, the underlying mechanisms contributing to auditory hypersensitivity are not well understood.

The acoustic startle response (ASR) is a protective response which usually manifests as a whole-body reflex following a sudden and unexpected stimulus [18]. Notably, the magnitude of the ASR is attenuated when a weak and non-startling auditory stimulus (prepulse) precedes an intense and startling stimulus, a phenomenon known as prepulse inhibition of acoustic startle response (PPI) [19]. PPI is commonly used as a behavioral paradigm of sensorimotor gating which is a normal physiological process responsible for sensory filtration. Impairments in sensorimotor gating can contribute to auditory hypersensitivity [6]. Frankland et al observed robustly decreased PPI and increased ASR in 8-to 17-year-old males with FXS [20]. Another study by Hessl et al also observed decreased PPI, although ASR was not reported, in 8-to 40-year-old male and female FXS individuals [21].

Previous studies in the FXS mouse model regarding PPI and ASR were variable. Some studies noted decreased PPI and ASR in young (post-natal day 23-25, P23-25) *FMR1*-KO mice [6] and 8-week old *FMR1*-KO mice [22]. Other studies observed increased PPI and reduced ASR in 8 to 12 weeks old *FMR1*-KO mice [20]. Similarly, increased PPI was observed in both 2-to 3-month-old *FMR1*-KO mice [23] and 7 to 10 weeks old *FMR1*-KO mice [24]. Chen et al also reported decreased ASR in the *FMR1*-KO mice [24]. However, these findings particularly support the involvement of the IC in auditory hypersensitivity, since the acoustic PPI pathway is believed to be mediated via the IC. Interestingly, this is consistent with previous studies which observed a complete disruption of acoustic PPI following lesions at the IC [25, 26].

The gap-induced inhibition of the acoustic startle (GPIAS) is another variant of the PPI paradigm, which involves using a gap in a continuous background noise as prepulse instead of an acoustic stimulus, and is mediated by the auditory cortex (AC) [27]. One study by McCullagh et al observed decreased GPIAS in mature (P55-P167) male and female *FMR1-*KO mice [28], indicating that auditory processing was impaired at the AC. Intriguingly, this aligns with previous observation of enhanced AC neuronal responses to auditory stimuli in male *FMR1*-KO mice, particularly during early development (P21) [29]. Together with heightened susceptibility for audiogenic seizure (AGS) expression between P20 and P30 in the *FMR1*-KO mice, this also suggests that auditory hypersensitivity can also be observed in young mice [17, 30]. Therefore, further investigation is required to understand whether early GPIAS deficits can possibly corroborate previous physiology studies and the role of the AC in auditory hypersensitivity.

Habituation, or reduction in ASR magnitude in response to repeating stimulus, is also an important sensory filtering mechanism [31] and atypical habituation is a common phenotype in both ASD and FXS [32, 33]. In the present study, we investigated both PPI using the GPIAS paradigm and habituation in the *FMR1-*KO mice at P15, P20 and P30. We observed no significant genotype differences in GPIAS at any of the developmental timepoints that we tested. However, significant sex and developmental difference were observed in the response latency of the ASR. Moreover, we also observed significant genotype, sex and developmental differences in the response duration of ASR. Finally, we also observed significant impairments in habituation and sensitization in the KO mice.

## Methods

### Animals

Wildtype (FVB.129P2-Pde6b+ Tyrc-ch/AntJ, Jax stock No: 004828) and *FMR1-*knockout (KO) (FVB.129P2-*Pde6b*^+^ *Tyr*^*c-ch*^ *Fmr1*^*tm1Cgr*^/J, Jax stock No: 004624) were obtained from the Jackson Laboratory (ME, USA) and maintained in the mouse housing facility at the Health Sciences Animal Resources Centre, University of Calgary. Overall, a total of 40 mice were tested, male WT (n=10), female WT (n=10), male KO (n=10) and female KO (n=10), with mouse pups first being tested at post-natal day 15 (P15), following which mice pups were returned to the dam and subsequently weaned and tested at P20 and group-housed (up to five mice per cage) with same sex littermates for further testing at P30. In the present study, each of our testing sessions was set several days apart to limit the carryover effect of previous auditory sessions. This is consistent with a previous study where water-restricted mice were trained to perform an auditory delayed match-to-sample task, wherein mice had to remember the auditory stimulus following a brief delay (1.5 s). In general, the behavioral performance was observed to decrease meaning that the mice did not recall the initial tone [34], with increasing delay duration (from 2 to 14 s).

Mouse housing cages were maintained at the animal facility with automated 12-hr light and dark cycle and access to food (standard mouse chow) and water ad libitum. Behavioral recordings, which included gap-induced inhibition of the acoustic startle (GPIAS) and acoustic startle response (ASR) were carried out between 9:00h and 19:00h. Furthermore, all recordings were performed in accordance with the guidelines outlined by the Canadian Council for Animal Care and approved by the Health Sciences Animal Care Committee at the University of Calgary.

### Apparatus for ASR and GPIAS

Acoustic startle tests with mice were conducted in a custom-made animal housing within a dark sound-proof chamber. The animal housing was fitted with four piezoelectric transducers which were responsible for converting animal movement into voltage signals. Stimulus parameters were designed using RPvds software (Tucker-Davis Technologies, Inc., Gainesville, FL, USA). Acoustic stimuli were delivered by BrainWare software (Tucker-Davis Technologies, Inc., Gainesville, FL, USA) via the RZ6 processor, and output signals from the animal housing were recorded and sampled at 25 kHz by the RZ5 processor (Tucker-Davis Technologies, Inc., Gainesville, FL, USA) and the BrainWare software. Two loudspeakers (MF1, Tucker-Davis Technologies, Gainesville, FL, USA), one for delivering the startle stimulus (20 ms, white noise, 106 dB SPL) and another for delivering a white background noise (70 dB SPL) and the prepulse gap (20 ms), were placed 10 cm from the animal housing. Prior to recording, the tone amplitude (expressed as dB SPL, re. 20 μPa) of both loudspeakers was calibrated at the same position using a condenser microphone (Model 2520, Larson-Davis Laboratories, USA) and a microphone preamplifier (Model 2200C, Larson-Davis Laboratories, USA).

### Procedure for ASR and GPIAS

Mice were allowed to acclimate to the testing chamber for approximately 10 minutes before testing. Following the acclimation period, a series of 20 startle stimulus were provided, which were later analyzed for habituation and sensitization. Another ten-minute break was provided before the GPIAS trials. The GPIAS trials consisted of a gap in the continuous background noise (prepulse) and the startle stimulus with an inter-stimulus interval (ISI) of 50 or 100 ms. These ISI were chosen for this study based on a previous study that observed robust GPIAS at 50 ms and 100 ms [35]. Overall, 24 trials were provided randomly consisting of 8 gap trials with 50 ms (ISI), 8 gap trials with 100 ms (ISI) and 8 startle-alone trials, with inter-trial intervals ranging between 15 to 25 s (shown in **Fig. 1**).

**Fig. 1.**
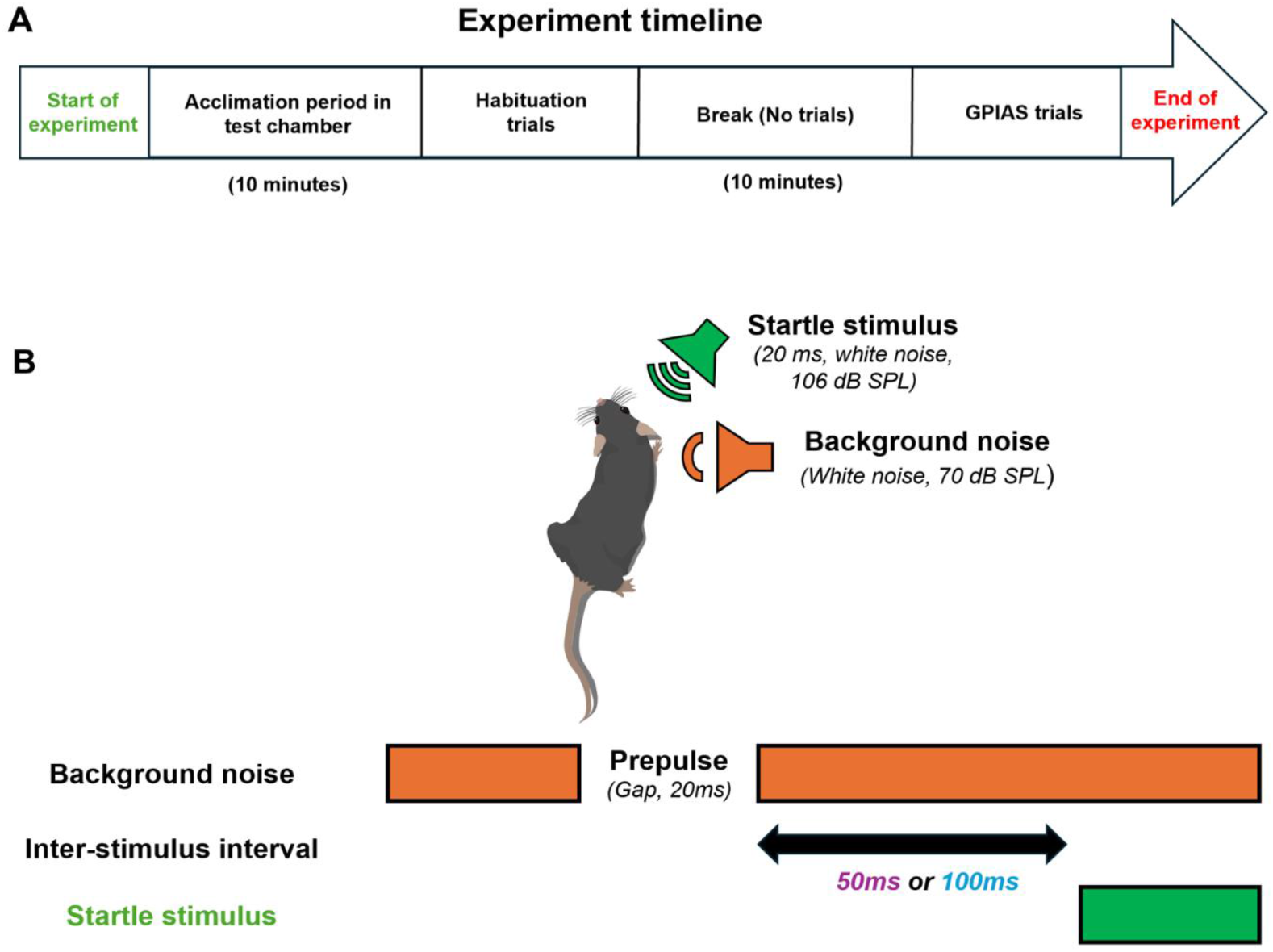
(A) Schematic diagram of the experiment timeline. (B) Methodology of GPIAS which includes a continuous background noise, gap in the background noise, startle stimulus and two GPIAS conditions at ISI 50 ms and 100 ms. Mouse figure adapted from *Scidraw, S. (2020). mouse top. Zenodo. https://doi.org/10.5281/zenodo.3925917 (Creative Commons 4.0 license)*

### Data processing

The original action potential data processed by BrainWare were stored as DAM files which were later processed with MATLAB. The following response properties were characterized based on the action potential data:

1. Amplitude of startle response: Magnitude of ASR was calculated as the root mean square (RMS) of the peaks within a fixed time window of 100 ms following stimulus onset.
2. Percentage of PPI: PPI was calculated by comparing the response magnitudes during prepulse trials with the startle-alone trials, shown by the following formula:

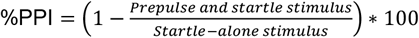
3. Latency: The latency of the startle response, i.e. is the time between stimulus onset and beginning of the startle response, was analyzed for all GPIAS conditions.
4. Response duration: Response duration was quantified by first analyzing the signal as 10 ms width bins. Baseline activity was established by calculating the root mean square (RMS) of consecutive 50 bins pre-stimulus (500 ms). RMS was further analyzed within 500 ms post-stimulus. A threshold was defined as the baseline mean and two times the standard deviation (mean ± 2SD). Th response duration was defined as the time from stimulus onset to the beginning of the first pair of consecutive bins with RMS value below the threshold, which indicated the offset of the response.
5. Habituation: Habituation score was calculated for each animal, to asses habituation [36], by analyzing the first eight trials from the habituation period that was performed prior to the GPIAS trials. Increased habituation scores would indicate that the mice’s startle response remained relatively large across repetitive trials, reflecting reduced habituation. Conversely, reduced scores indicate a reduction in response magnitude across repetitive trials, reflecting enhanced habituation.

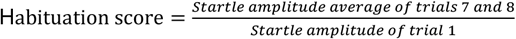
6. Sensitization: The sensitization score was calculated to assess sensitization [37] by comparing the subsequent three consecutive trials with the first trial during the habituation period. Increased sensitization scores would indicate that the startle response magnitude increased between the first stimulus presentation and subsequent stimuli, and vice-versa.

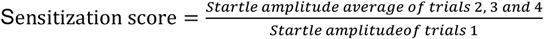

### Statistics

GraphPad Prism 10.2.3 (GraphPad Software, San Diego, California) was used to perform all statistical tests and produce the figures. All data are shown as Mean ± SEM and α was set at 0.05. For all figures, a three-way repeated measure ANOVA was performed (since the same mice were tested at different developmental timepoints) along with post-hoc Fisher’s test. Non-parametric Spearman’s correlation analysis was also performed to investigate whether the startle response magnitude was dependent on mice’s weight.

## Results

### Startle response magnitude increased with development

We first analyzed the acoustic startle response (ASR) from startle-alone trials during the gap-induced inhibition of the acoustic startle (GPIAS) period, and quantified mice weight to investigate whether there was a correlation between mice weight and startle response. There was a non-significant trend for genotype difference in startle response magnitude (p= 0.0673), with post-hoc analysis showing decreased ASR magnitude in the female *FMR1-*knockout (KO) mice compared to their wild-type (WT) counterparts at P30 (p= 0.0102) (shown in **Fig. 2A**). We also observed no significant differences between the weight of KO and WT mice at all ages (p= 0.9032) (shown in **Fig. 2B**). There was significant sex difference in startle response (p= 0.0004), particularly at P30 between the male and female KO (p= 0.0014). Significant sex difference was also observed within mice weight (p= 0.0035), and further post-hoc analysis showed significant difference at P30 between male and female KO (p= 0.0013). In terms of development, the magnitude of the startle response (p= <0.0001) and mice weight (p= <0.0001) increased with maturation, with post-hoc analysis showing several group differences which are included in the supplementary tables **(Table S1 and S2)**.

**Fig. 2.**
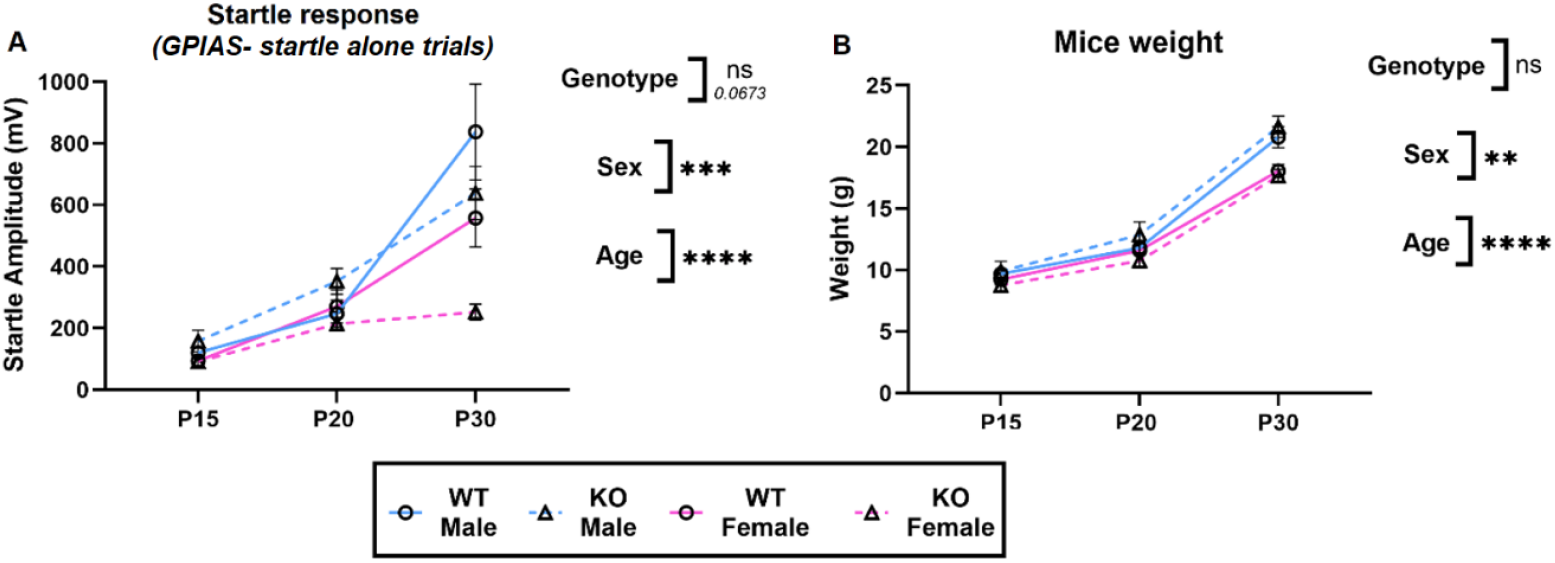
Startle response magnitude and mice weight at different developmental ages in both the female and male *FMR1*-KO and WT mice. (A) Comparison of startle response magnitude during early and late development. (B) Comparison of mice weight during early and late development. Three-way RM ANOVA was performed along with post-hoc Fisher’s test (**: 0.0021, ***:0.0002, ****: <0.0001).

Furthermore, we observed that there was no significant correlation between startle response magnitude and body weight in both the female WT and female KO groups at all the developmental timepoints. However, in males some significant correlations were observed in male WT at P15 and P20, and in male KO at P20 and P30 **(Table S3)**, suggesting that while startle response is independent of the body weight in the female group, the weight of the mice may affect the startle response in the male group.

### No significant genotype difference in gap-induced prepulse inhibition during early or late development

We also quantified GPIAS in both male and female *FMR1-*KO and WT mice at different developmental timepoints. At both GPIAS conditions, there were no significant genotype (ISI: 50 ms, p= 0.9454; ISI: 100 ms, p= 0.9568) or sex difference (ISI: 50 ms, p= 0.7411; ISI: 100 ms, p= 0.9631) at P15, P20 and P30. However, in terms of developmental changes, we observed increasing GPIAS with maturation (ISI 50 ms, p= 0.0039) during early auditory development (shown in **Fig. 3A and B**). This was evident in further post-hoc analysis **(Table S4)** which showed that there were significant age differences between P15 and P30 WT male (p= 0.0087), P15 and P30 KO female (p= 0.0106) and between P20 and P30 KO female (p= 0.0022).

**Fig. 3.**
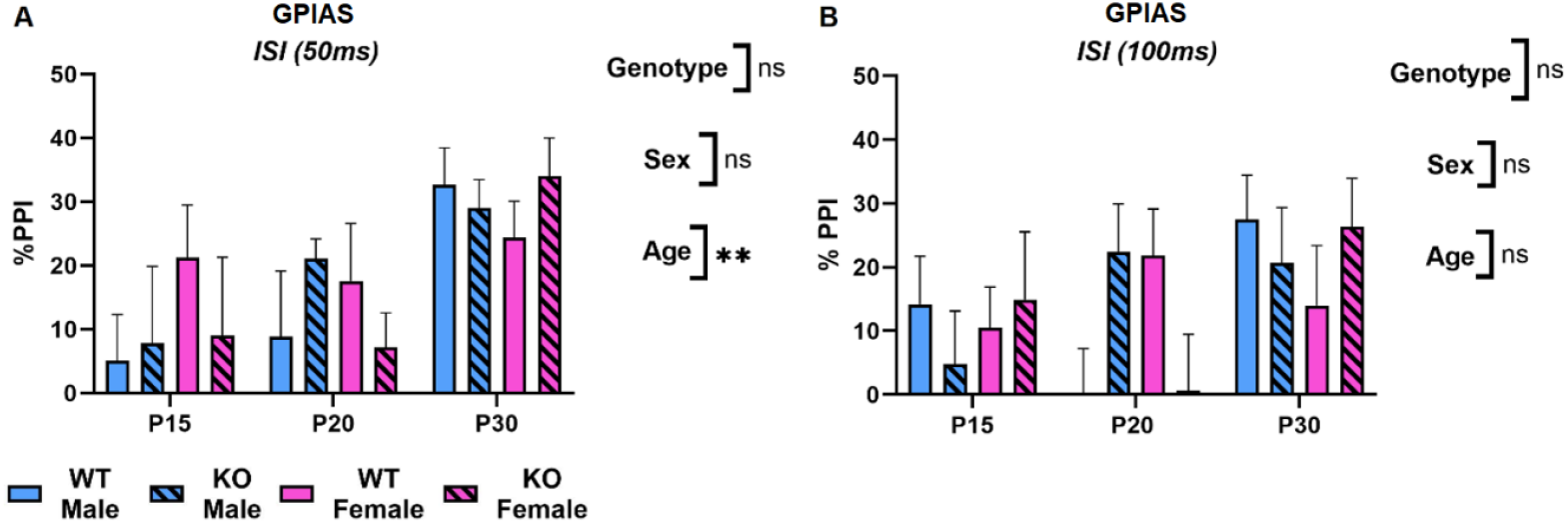
Development of PPI in both the female and male *FMR1*-KO and WT mice. (A) Comparison of PPI at 50 ms ISI during early and late development. (B) Comparison of PPI at 100 ms ISI during early and late development. Three-way RM ANOVA was performed along with the post-hoc Fisher’s test (**: 0.0021).

### No genotype difference but significant developmental change was observed in latency under all GPIAS conditions

We also analyzed latency of the ASR under various GPIAS conditions during development. We did not observe any significant genotype difference at any GPIAS condition (ISI 50 ms, p= 0.5979; ISI 100 ms, p= 0.5900; Startle-alone, p= 0.8764) (shown in **Fig. 4A-C**). In addition, we only observed significant sex difference in the startle-alone GPIAS condition (p= 0.0250) but further post-hoc Fisher’s analysis did not show any significant group difference. Finally, in terms of developmental change, there were significant differences at all GPIAS conditions (ISI 50 ms, p= <0.0001; ISI 100 ms, p= <0.0001; Startle-alone, p= 0.0040) with further post-hoc Fisher’s analysis showing significant age differences within several groups which are included in the supplementary tables **(Table S5-7)**.

**Fig. 4.**
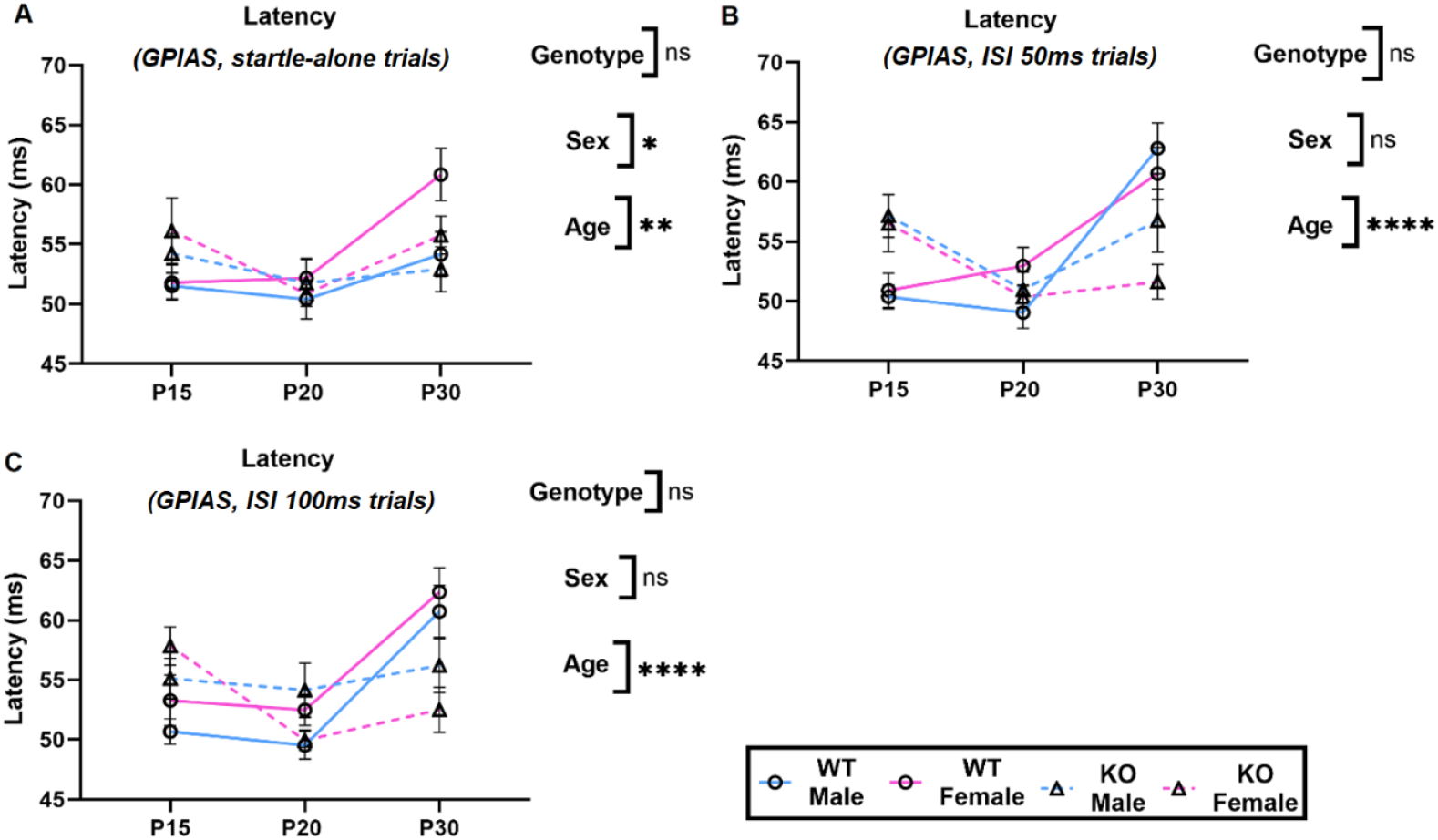
Latency of acoustic startle response during GPIAS trials at various developmental timepoints in the female and male *FMR1*-KO and WT mice. (A) Comparison of latency of startle response 50 ms (ISI) after prepulse at P15, P20 and P30. (B) Comparison of latency of startle response 100 ms (ISI) after prepulse at P15, P20 and P30. (C) Comparison of latency of startle response following no prepulse at P15, P20 and P30. Three-way RM ANOVA was performed along with post-hoc Fisher’s test (*:0.0332, **: 0.0021, ***:0.0002, ****: <0.0001).

### Significant genotype, sex and developmental difference in response duration of the acoustic startle response

The final response parameter that we analyzed from the GPIAS trials was response duration of the ASR (shown in **Fig. 5 A-C**). We observed that there were significant genotype differences in response duration (ISI 50 ms, p= 0.0397; ISI 100 ms, p= 0.0269), and further post-hoc Fisher’s analysis indicated significantly increased response duration in P20 male KO mice (ISI 50 ms, p= 0.0145) and P15 male KO mice (ISI 100 ms, p= 0.0058) compared to their respective age-matched male WT mice.. In terms of sex, we observed an overall significant difference within all three GPIAS conditions (ISI 50 ms, p= 0.0007; ISI 100 ms, p= 0.0085; Startle-alone, p= 0.0038), with post-hoc analysis showing significantly decreased response durations in P30 female KO mice (ISI 100 ms, p= 0.0416) and P30 female WT mice (Startle-alone, p=0.0379) compared to their male counterparts.

**Fig. 5.**
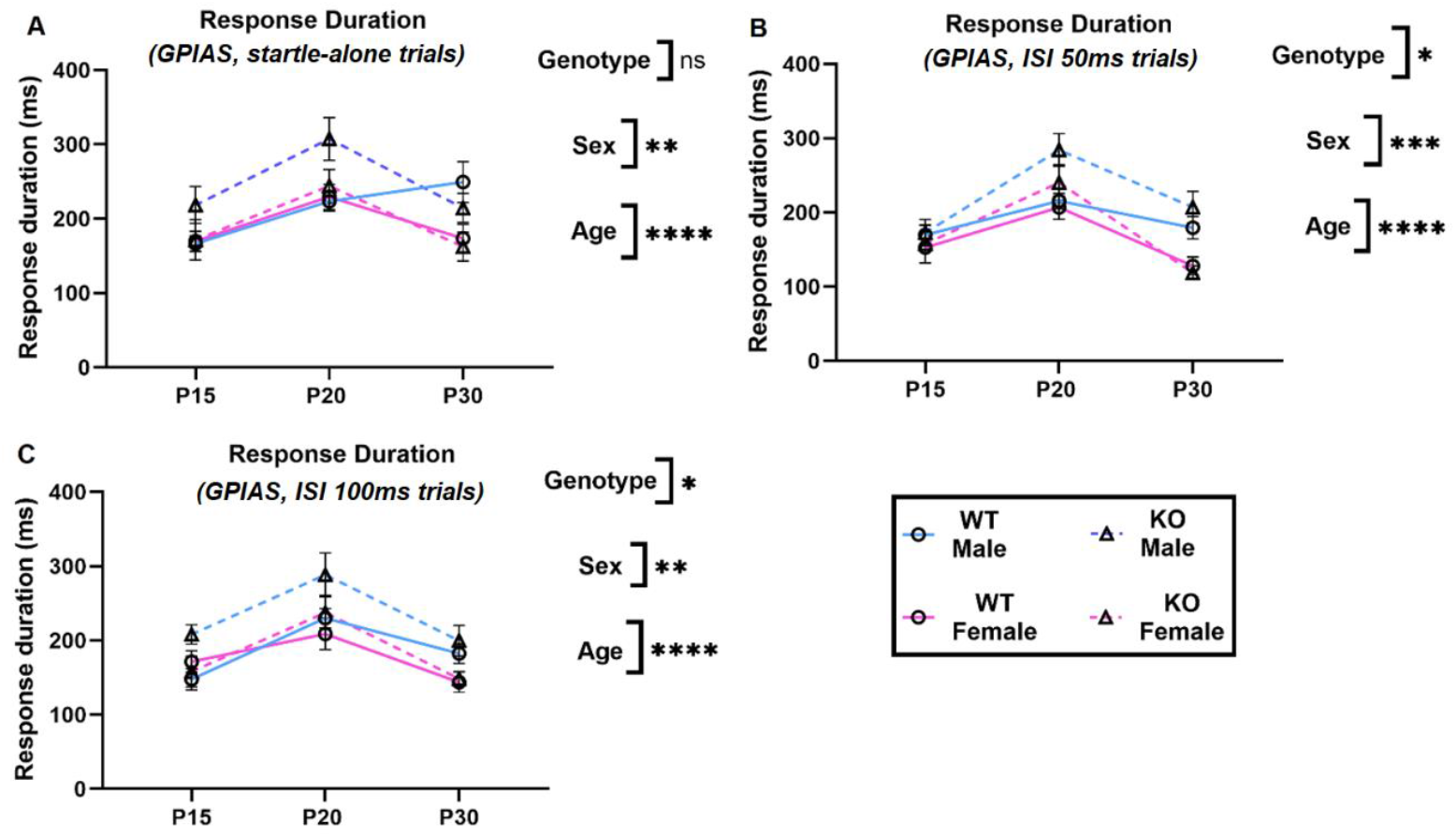
Response duration of acoustic startle response during GPIAS trials at various developmental timepoints in the female and male *FMR1*-KO and WT mice. (A) Comparison of response duration of startle response 50 ms (ISI) after prepulse at P15, P20 and P30. (B) Comparison of response duration of startle response 100 ms (ISI) after prepulse at P15, P20 and P30. (C) Comparison of response duration of startle response following no prepulse at P15, P20 and P30. Three-way RM ANOVA was performed along with post-hoc Fisher’s test (*:0.0332, **: 0.0021, ***:0.0002, ****: <0.0001).

Moreover, we also observed significant developmental differences in response duration at all the GPIAS conditions (p= <0.0001), with post-hoc analysis showing that some groups demonstrated a significant increase in response duration between P15 and P20, while other groups displayed a decrease in response duration between P20 and P30. Our post-hoc analysis has also been included in the supplementary tables **(Table S8-10)**.

### Significant genotype differences are observed in both habituation and sensitization during early and late development

Finally, we also analyzed both habituation and sensitization during the habituation period prior to the GPIAS trials in our study. Significant genotype differences were observed in habituation (p= 0.0278) (shown in **Fig. 6A**), although post-hoc Fisher’s analysis only noted a non-significant trend (p= 0.0742) for increased habituation score, or reduced habituation following consecutive trials, in the KO females compared to their WT counterparts at P30. In addition, significant genotype differences were observed in sensitization (p= 0.0050) (shown in **Fig. 6B**), and further post-hoc Fisher’s analysis showed that the P15 KO female mice displayed higher sensitization scores compared to the female WT mice (p= 0.0331). No significant sex difference was observed in habituation (p= 0.5101) or sensitization (p= 0.4535). We also did not observe any significant age difference in habituation (p= 0.9262) or sensitization (p= 0.1776).

**Fig. 6.**
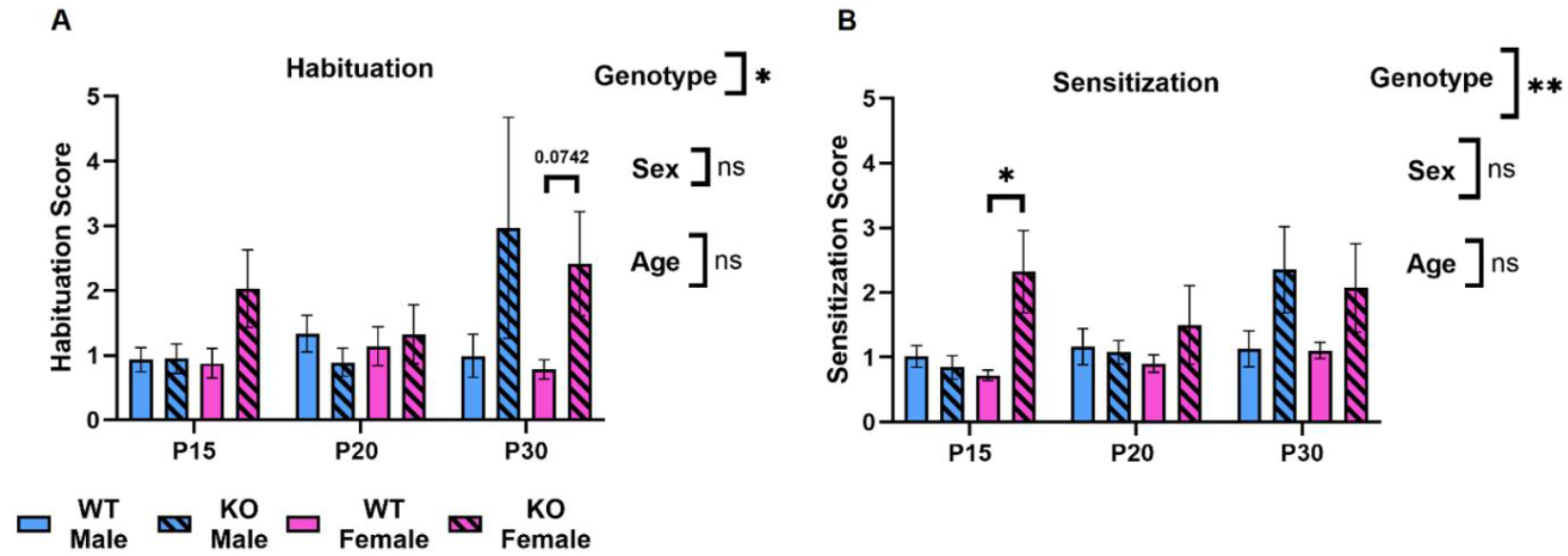
Habituation and sensitization during early and late auditory development in the female and male *FMR1*-KO and WT mice. (A) Comparison of habituation at P15, P20 and P30. (B) Comparison of sensitization at P15, P20 and P30. Three-way RM ANOVA was performed along with post-hoc Fisher’s test (*:0.0332, **: 0.0021, ***:0.0002, ****: <0.0001).

## Discussion

We utilized a behavioral paradigm called prepulse inhibition (PPI), particularly gap-induced inhibition of the acoustic startle (GPIAS) to investigate whether impairments in sensorimotor gating can contribute to auditory hypersensitivity during early development in a mouse model of Fragile X Syndrome (FXS). Our results indicated there were no significant genotype differences in GPIAS, magnitude of acoustic startle response (ASR) or response latency, but significant genotype differences were observed in duration of the startle response. Moreover, significant sex differences were also observed in startle response magnitude and latency. Finally, we also observed significant genotype differences in both habituation and sensitization, which could be a contributing factor to auditory hypersensitivity during early auditory development.

In terms of developmental changes at post-natal days 15 (P15), 20 (P20) and 30 (P30), there were significant age differences in startle response magnitude, latency, and duration, indicating that the acoustic response properties change with development irrespective of genotypes and sexes.

### *FMR1-*KO mice displayed no significant genotype difference in gap-induced prepulse inhibition and startle response magnitude during development

Here, we investigated whether PPI deficits can be observed during early development (P15, P20 and P30) using the GPIAS paradigm (with interstimulus intervals 50 ms and 100 ms) in the *FMR1-*KO mice. Our results indicate that although there was a trend, no significant genotype differences in startle response magnitude (shown in **Fig. 2A**) was observed, which is similar to findings from another study in FXS individuals [20].

Previously, one study by Yun et al observed that the Fragile X Messenger Ribonucleoprotein (FMRP) expression in the pedunculopontine tegmental nucleus (PPTg), which receives direct auditory projections from the auditory cortex (AC), decreased after post-natal 3-4 weeks of age in the WT mice. Yun et al also observed that with maturation the ASR magnitude remained the same in the KO mice unlike the WT mice, indicating that in the absence of FMRP further development of ASR was arrested in the KO mice [38]. This could explain the trend that we observed towards reduced ASR magnitude in the KO female mice compared to their WT counterparts at P30 (shown in **Fig. 2A**).

Furthermore, we also did not observe any significant genotype difference in GPIAS during early development (shown in **Fig. 3**). One explanation for the absence of significant difference in GPIAS between the *FMR1-*KO and WT mice could be attributed to the age of the mice tested in this study. Our results indicated that there was an overall effect of age in the GPIAS test (ISI: 50 ms), indicating that GPIAS improved with age. This is consistent with one study which observed that improved GPIAS could be observed between ages 4 and 8 weeks in WT mice, possibly due to a developmental decrease in dendritic spine number at the AC during this period [39]. The AC has previously been suggested to play an important role in GPIAS, since bilateral lesions at the AC disrupted GPIAS but not auditory PPI [40]. Overall, our results suggest that GPIAS deficits in the *FMR1-*KO mice do not appear early in development and may be more reliably observed in mature mice.

### Significant developmental changes were observed in startle response magnitude

In the present study we also characterized the early development of the startle response magnitude following the startle stimulus. The primary neural pathway responsible for mediating the startle response originates from the spiral ganglion cells of the cochlea, which sends auditory inputs to the cochlear root neurons (CRN). The CRN innervates neurons in the caudal pontine reticular nucleus (PnC), which in turn projects to the spinal motoneurons causing rapid constriction of the skeletal muscles [27]. In terms of development, ASR was observed to emerge in rodents around P14 [41], which aligned with the period during which the inner hair cells (IHC) of the cochlea reached full maturation while the outer hair cells (OHC) developed further until P28 [42]. In one study, Pietropaolo et al noted that the magnitude of the startle response increased with maturation between 4 and 8 weeks in the C57Bl/6 mice [43]. Another study also observed that the ASR was first detectable at post-natal week 2 and the magnitude of the ASR gradually increased until 8 weeks of age in WT mice [38]. These findings are consistent with our current results wherein the startle response magnitude increased between P15, P20 and P30 (shown in **Fig. 2A**).

In addition, our findings also showed no genotype difference in body weight and that the ASR magnitude was not always correlated with weight. Overall, this suggests that while body weight may influence the ASR magnitude during development, increasing ASR magnitude could mainly reflect the early development period of the OHC and IHC in the cochlea, which are important for modulating ASR amplitude [44], within the ASR pathway.

### Aging led to significant changes in latency of acoustic startle response

Another response parameter that we characterized within the GPIAS trials was response latency of ASR. We observed an increase in startle latency with age at all GPIAS conditions (shown in **Fig. 4**). Increasing startle latency with age could be attributed to delayed neural processing. Particularly, one study by Tresch et al observed delayed startle stimuli evoked movement of the sternocleidomastoid (SCM) muscle in humans, possibly due to neural processing delays at the brainstem, in older individuals (age 70 ± 11) compared to young adults (age 24 ± 1) [45]. However, given that the latency of ASR has not been widely studied in mice, further investigation is required to understand the mechanisms underlying the maturation of ASR latency.

### Significant genotype and developmental difference were observed in response duration

We also characterized the response duration of the acoustic startle response within the GPIAS trials. We observed significant genotype differences (GPIAS trial; ISI 50 ms and 100 ms) and developmental changes under all three GPIAS conditions (shown in **Fig. 5**). Particularly, our findings indicate that the response duration initially increased between P15 and P20, followed by a decline between P20 and P30. Although the literature on the response duration of the startle response is limited, the developmental changes in response duration could reflect the maturation of the auditory pathway involved in the startle response.

FMRP is required for regulating neural excitability and synaptic transmission via interactions with several ion channels that include the β4 subunit of large voltage and calcium dependent potassium channels called BK channel. In the absence of FMRP, the activity of the BK channel is reduced. This leads to long action potential duration, enhanced calcium influx into the presynaptic terminal, elevated neurotransmitter release and defects in short-term plasticity [46]. Normal BK channel function is required for preferentially modulating excitatory glutaminergic neurotransmitter release [47]. Therefore, BK channel dysfunction could lead to disrupted excitatory-inhibitory balance, which was previously noted in FXS [48]. Consequently, increased excitability of neurons could prolong neural firing and response duration as observed in our study. Overall, our observation of significant genotype and developmental differences in duration of the ASR with the *FMR1*-KO mouse model is a novel finding that has not been previously reported with the GPIAS paradigm, and further studies are required to understand whether atypical response duration can be observed at the auditory structures involved within the ASR and GPIAS pathway.

### Significant genotype differences were observed in habituation and sensitization

Previously, Nielsen et al noted an absence of short-term habituation in the *FMR1*-KO mice (12-15 weeks old male) in response to repeated acoustic stimulus (110 dB or 120 dB) [49]. Similarly, previous studies with EEG/ERP also observed impairments in habituation of the N1 component in both individuals with FXS and the *FMR1*-KO mouse model [50, 51]. Therefore, we also characterized habituation which is the decrease in ASR magnitude over a series of trials. Our findings indicate that there were significant genotype differences in habituation, indicating that the *FMR1*-KO mice displayed decreased habituation over consecutive trials (shown in **Fig. 6A**).

Besides habituation, we also analyzed sensitization which is the increase in ASR magnitude in consecutive trials following the first trial. Madsen et al previously noted significantly increased sensitization in boys and girls (ages between 8 and 12 years old) diagnosed with ASD compared to the control group [52]. Another study by Möhrle et al also observed increased sensitization in both male and female *Cntnap2*-KO rats, an animal model of ASD, compared to WT rats [37]. Although, sensitization to auditory stimulus remains largely unexplored in FXS, some studies observed increased sensitization to visual stimulus in individuals with FXS [53, 54]. Consistent with the previous studies, we also noted increased sensitization in KO mice compared with WT mice (shown in **Fig. 6B**). Together, these data suggest that the KO mice may display disruptions in their ability to adapt to repeating stimuli.

Activation of BK channels was also previously implicated to play a role in the reduction of the synaptic strength at the PnC, leading to short-term habituation following consecutive stimulus [31]. Therefore, a possible explanation for decreased auditory short-term habituation in the KO mice could be attributed to reduced BK channel function at the PnC in the absence of FMRP. Sensitization, on the other hand is believed to be mediated by the amygdala [55], based on previous studies which observed reduced sensitization of startle response following lesions or blockade of the amygdala [56, 57]. Interestingly, absence of FMRP also leads to the dysfunction of the amygdala [58], which can possibly lead to increased sensitization in the *FMR1-*KO mice compared to the WT mice.

## Conclusion

Auditory hypersensitivity is a common and debilitating phenotype of FXS. In the present study, we observed significant changes in behavioral response properties with development and differences between females and males which is an important consideration for future study design. We also observed significant changes between WT and FXS mouse models in habituation during early development, which could be an ideal window for addressing auditory hypersensitivity in FXS. Further, future mechanistic studies could provide more insight into how individuals may adapt to the environment differently in FXS.

## Acknowledgements

We would like to acknowledge Dorit Möhrle for providing the MATLAB code for data export.

## Statement of Ethics

We have reviewed and abided by the statement of ethical standards for manuscripts submitted to Developmental Neuroscience. All procedures in this study were performed in accordance with the recommendations in the Canadian Council for Animal Care. The protocol of this study was approved by the Health Sciences Animal Care Committee of the University of Calgary and could be provided upon request. All efforts were made to minimize the numbers of animals used and their suffering.

## Author Contributions

Abdullah Abdullah participated in designing the experiment, recording, data analysis and interpretation, writing and revising the manuscript. Xiuping Liu participated in designing experiments. Kartikeya Murari contributed to data analysis and interpretation. Jun Yan participated in designing experiments, data analysis and interpretation, and editing and revising the manuscript. Ning Cheng designed the study, participated in data analysis and interpretation, and revised the manuscript.

## Data Availability Statement

Data will be made available on request.

