## Supplementary Table for "Early developmental characterization of gap-induced prepulse inhibition and habituation in the Fragile X Syndrome mouse model"

### Supplementary Material

**Table S1: Pair-wise pose hoc Fisher's group comparison following three-way RM ANOVA for startle response (PPI- startle alone trials).** Values in green indicate significant age difference and values in red indicate significant sex difference.

| Parameter | Group comparisons | Summary of significance |
|---|---|---|
| Startle response (PPI- startle alone trials) | P15 Male WT vs P15 Male KO | **n.s** |
| | P15 Male WT vs P15 Female WT | **n.s** |
| | P15 Male WT vs P20 Male WT | **0.0076** |
| | P15 Male WT vs P30 Male WT | **0.0017** |
| | P15 Male KO vs P15 Female KO | **n.s** |
| | P15 Male KO vs P20 Male KO | **<0.0001** |
| | P15 Male KO vs P30 Male KO | **<0.0001** |
| | P15 Female WT vs P15 Female KO | **n.s** |
| | P15 Female WT vs P20 Female WT | **0.0049** |
| | P15 Female WT vs P30 Female WT | **n.s** |
| | P15 Female KO vs P20 Female KO | **<0.0001** |
| | P15 Female KO vs P30 Female KO | **<0.0001** |
| | P20 Male WT vs P20 Male KO | **n.s** |
| | P20 Male WT vs P20 Female WT | **n.s** |
| | P20 Male WT vs P30 Male WT | **0.0108** |
| | P20 Male KO vs P20 Female KO | **n.s** |
| | P20 Male KO vs P30 Male KO | **0.0007** |
| | P20 Female WT vs P20 Female KO | **n.s** |
| | P20 Female WT vs P30 Female WT | **n.s** |
| | P20 Female KO vs P30 Female KO | **n.s** |
| | P30 Male WT vs P30 Male KO | **n.s** |
| | P30 Male WT vs P30 Female WT | **n.s** |
| | P30 Male KO vs P30 Female KO | **0.0014** |
| | P30 Female WT vs P30 Female KO | **n.s** |

**Table S2: Pair-wise post-hoc Fisher's group comparison following three-way RM ANOVA for mice body weight.** Values in green indicate significant age difference and values in red indicate significant sex difference.

| Parameter | Group comparisons | Summary of significance |
|---|---|---|
| **Mice body weight** | P15 Male WT vs P15 Male KO | **n.s** |
| | P15 Male WT vs P15 Female WT | **n.s** |
| | P15 Male WT vs P20 Male WT | **<0.0001** |
| | P15 Male WT vs P30 Male WT | **<0.0001** |
| | P15 Male KO vs P15 Female KO | **n.s** |
| | P15 Male KO vs P20 Male KO | **<0.0001** |
| | P15 Male KO vs P30 Male KO | **<0.0001** |
| | P15 Female WT vs P15 Female KO | **n.s** |
| | P15 Female WT vs P20 Female WT | **<0.0001** |
| | P15 Female WT vs P30 Female WT | **n.s** |
| | P15 Female KO vs P20 Female KO | **n.s** |
| | P15 Female KO vs P30 Female KO | **n.s** |
| | P20 Male WT vs P20 Male KO | **n.s** |
| | P20 Male WT vs P20 Female WT | **n.s** |
| | P20 Male WT vs P30 Male WT | **<0.0001** |
| | P20 Male KO vs P20 Female KO | **n.s** |
| | P20 Male KO vs P30 Male KO | **<0.0001** |
| | P20 Female WT vs P20 Female KO | **n.s** |
| | P20 Female WT vs P30 Female WT | **n.s** |
| | P20 Female KO vs P30 Female KO | **n.s** |
| | P30 Male WT vs P30 Male KO | **n.s** |
| | P30 Male WT vs P30 Female WT | **n.s** |
| | P30 Male KO vs P30 Female KO | **0.0013** |
| | P30 Female WT vs P30 Female KO | **n.s** |

**Table S3: Non-parametric Spearman's correlation analysis between startle response (PPI- startle alone trials) and mice weight.** Values highlighted in blue in each column are the correlation co-efficient (r) values with weak-moderate correlations in bold. Unhighlighted values in each column are the p-values with non-significant values in red.

| Groups | | Age | | |
|---|---|---|---|---|
| | | P15 | P20 | P30 |
| Male WT | r | 0.7333 | 0.7939 | **-0.3455** |
| | p | 0.0202 | 0.0088 | 0.3304 |
| Male KO | r | **0.5030** | 0.6606 | 0.8303 |
| | p | 0.1440 | 0.0438 | 0.0047 |
| Female WT | r | **0.3697** | **0.1152** | **0.2310** |
| | p | 0.2957 | 0.7589 | 0.5190 |
| Female KO | r | **0.2000** | **0.3455** | **0.1758** |
| | p | 0.5837 | 0.3304 | 0.6321 |

**Table S4: Pair-wise pose hoc Fisher's group comparison following three-way RM ANOVA for PPI (ISI 50ms trials).** Values in green indicate significant age difference.

| Parameter | Group comparisons | Summary of significance |
|---|---|---|
| PPI (ISI 50ms trials) | P15 Male WT vs P15 Male KO | n.s |
| | P15 Male WT vs P15 Female WT | n.s |
| | P15 Male WT vs P20 Male WT | n.s |
| | P15 Male WT vs P30 Male WT | 0.0087 |
| | P15 Male KO vs P15 Female KO | n.s |
| | P15 Male KO vs P20 Male KO | n.s |
| | P15 Male KO vs P30 Male KO | n.s |
| | P15 Female WT vs P15 Female KO | n.s |
| | P15 Female WT vs P20 Female WT | n.s |
| | P15 Female WT vs P30 Female WT | n.s |
| | P15 Female KO vs P20 Female KO | n.s |
| | P15 Female KO vs P30 Female KO | 0.0106 |
| | P20 Male WT vs P20 Male KO | n.s |
| | P20 Male WT vs P20 Female WT | n.s |
| | P20 Male WT vs P30 Male WT | n.s |
| | P20 Male KO vs P20 Female KO | n.s |
| | P20 Male KO vs P30 Male KO | n.s |
| | P20 Female WT vs P20 Female KO | n.s |
| | P20 Female WT vs P30 Female WT | n.s |
| | P20 Female KO vs P30 Female KO | 0.0022 |
| | P30 Male WT vs P30 Male KO | n.s |
| | P30 Male WT vs P30 Female WT | n.s |
| | P30 Male KO vs P30 Female KO | n.s |
| | P30 Female WT vs P30 Female KO | n.s |

**Table S5: Pair-wise post-hoc Fisher's group comparison following three-way RM ANOVA for latency (PPI- ISI 50ms trials).** Values in green indicate significant age difference.

| Parameter | Group comparisons | Summary of significance |
|---|---|---|
| **Latency (PPI- ISI 50ms trials)** | P15 Male WT vs P15 Male KO | **n.s** |
| | P15 Male WT vs P15 Female WT | **n.s** |
| | P15 Male WT vs P20 Male WT | **n.s** |
| | P15 Male WT vs P30 Male WT | **0.0007** |
| | P15 Male KO vs P15 Female KO | **n.s** |
| | P15 Male KO vs P20 Male KO | **n.s** |
| | P15 Male KO vs P30 Male KO | **n.s** |
| | P15 Female WT vs P15 Female KO | **n.s** |
| | P15 Female WT vs P20 Female WT | **n.s** |
| | P15 Female WT vs P30 Female WT | **0.0096** |
| | P15 Female KO vs P20 Female KO | **0.0314** |
| | P15 Female KO vs P30 Female KO | **n.s** |
| | P20 Male WT vs P20 Male KO | **n.s** |
| | P20 Male WT vs P20 Female WT | **n.s** |
| | P20 Male WT vs P30 Male WT | **<0.0001** |
| | P20 Male KO vs P20 Female KO | **n.s** |
| | P20 Male KO vs P30 Male KO | **n.s** |
| | P20 Female WT vs P20 Female KO | **n.s** |
| | P20 Female WT vs P30 Female WT | **n.s** |
| | P20 Female KO vs P30 Female KO | **n.s** |
| | P30 Male WT vs P30 Male KO | **n.s** |
| | P30 Male WT vs P30 Female WT | **n.s** |
| | P30 Male KO vs P30 Female KO | **n.s** |
| | P30 Female WT vs P30 Female KO | **n.s** |

**Table S6: Pair-wise post-hoc Fisher's group comparison following three-way RM ANOVA for latency (PPI- ISI 100ms trials).** Values in green indicate significant age difference.

| Parameter | Group comparisons | Summary of significance |
|---|---|---|
| Latency (PPI- ISI 100ms trials) | P15 Male WT vs P15 Male KO | n.s |
| | P15 Male WT vs P15 Female WT | n.s |
| | P15 Male WT vs P20 Male WT | n.s |
| | P15 Male WT vs P30 Male WT | 0.0032 |
| | P15 Male KO vs P15 Female KO | n.s |
| | P15 Male KO vs P20 Male KO | n.s |
| | P15 Male KO vs P30 Male KO | n.s |
| | P15 Female WT vs P15 Female KO | n.s |
| | P15 Female WT vs P20 Female WT | n.s |
| | P15 Female WT vs P30 Female WT | 0.0033 |
| | P15 Female KO vs P20 Female KO | 0.0044 |
| | P15 Female KO vs P30 Female KO | n.s |
| | P20 Male WT vs P20 Male KO | n.s |
| | P20 Male WT vs P20 Female WT | n.s |
| | P20 Male WT vs P30 Male WT | 0.0002 |
| | P20 Male KO vs P20 Female KO | n.s |
| | P20 Male KO vs P30 Male KO | n.s |
| | P20 Female WT vs P20 Female KO | n.s |
| | P20 Female WT vs P30 Female WT | n.s |
| | P20 Female KO vs P30 Female KO | n.s |
| | P30 Male WT vs P30 Male KO | n.s |
| | P30 Male WT vs P30 Female WT | n.s |
| | P30 Male KO vs P30 Female KO | n.s |
| | P30 Female WT vs P30 Female KO | n.s |

**Table S7: Pair-wise post-hoc Fisher's group comparison following three-way RM ANOVA for latency (PPI- startle alone trials).** Values in green indicate significant age difference.

| Parameter | Group comparisons | Summary of significance |
|---|---|---|
| **Latency (PPI- startle alone trials)** | P15 Male WT vs P15 Male KO | **n.s** |
| | P15 Male WT vs P15 Female WT | **n.s** |
| | P15 Male WT vs P20 Male WT | **n.s** |
| | P15 Male WT vs P30 Male WT | **n.s** |
| | P15 Male KO vs P15 Female KO | **n.s** |
| | P15 Male KO vs P20 Male KO | **n.s** |
| | P15 Male KO vs P30 Male KO | **n.s** |
| | P15 Female WT vs P15 Female KO | **n.s** |
| | P15 Female WT vs P20 Female WT | **n.s** |
| | P15 Female WT vs P30 Female WT | **0.0076** |
| | P15 Female KO vs P20 Female KO | **n.s** |
| | P15 Female KO vs P30 Female KO | **n.s** |
| | P20 Male WT vs P20 Male KO | **n.s** |
| | P20 Male WT vs P20 Female WT | **n.s** |
| | P20 Male WT vs P30 Male WT | **n.s** |
| | P20 Male KO vs P20 Female KO | **n.s** |
| | P20 Male KO vs P30 Male KO | **n.s** |
| | P20 Female WT vs P20 Female KO | **n.s** |
| | P20 Female WT vs P30 Female WT | **0.0324** |
| | P20 Female KO vs P30 Female KO | **0.0118** |
| | P30 Male WT vs P30 Male KO | **n.s** |
| | P30 Male WT vs P30 Female WT | **n.s** |
| | P30 Male KO vs P30 Female KO | **n.s** |
| | P30 Female WT vs P30 Female KO | **n.s** |

**Table S8: Pair-wise post-hoc Fisher's group comparison following three-way RM ANOVA for response duration (PPI- ISI 50 ms trials).** Values in green indicate significant age difference.

| Parameter | Group comparisons | Summary of significance |
|---|---|---|
| Response duration (PPI- ISI 50ms trials) | P15 Male WT vs P15 Male KO | n.s |
| | P15 Male WT vs P15 Female WT | n.s |
| | P15 Male WT vs P20 Male WT | 0.0019 |
| | P15 Male WT vs P30 Male WT | n.s |
| | P15 Male KO vs P15 Female KO | n.s |
| | P15 Male KO vs P20 Male KO | 0.0063 |
| | P15 Male KO vs P30 Male KO | n.s |
| | P15 Female WT vs P15 Female KO | n.s |
| | P15 Female WT vs P20 Female WT | n.s |
| | P15 Female WT vs P30 Female WT | n.s |
| | P15 Female KO vs P20 Female KO | 0.0240 |
| | P15 Female KO vs P30 Female KO | n.s |
| | P20 Male WT vs P20 Male KO | 0.0145 |
| | P20 Male WT vs P20 Female WT | n.s |
| | P20 Male WT vs P30 Male WT | n.s |
| | P20 Male KO vs P20 Female KO | n.s |
| | P20 Male KO vs P30 Male KO | n.s |
| | P20 Female WT vs P20 Female KO | n.s |
| | P20 Female WT vs P30 Female WT | <0.0001 |
| | P20 Female KO vs P30 Female KO | 0.0010 |
| | P30 Male WT vs P30 Male KO | n.s |
| | P30 Male WT vs P30 Female WT | n.s |
| | P30 Male KO vs P30 Female KO | n.s |
| | P30 Female WT vs P30 Female KO | n.s |

**Table S9: Pair-wise post-hoc Fisher's group comparison following three-way RM ANOVA for response duration (PPI- ISI 100 ms trials).** Values in green indicate significant age difference and values in red indicate significant sex difference.

| Parameter | Group comparisons | Summary of significance |
|---|---|---|
| Response duration (PPI- ISI 100ms trials) | P15 Male WT vs P15 Male KO | **0.0058** |
| | P15 Male WT vs P15 Female WT | **n.s** |
| | P15 Male WT vs P20 Male WT | **0.0007** |
| | P15 Male WT vs P30 Male WT | **n.s** |
| | P15 Male KO vs P15 Female KO | **n.s** |
| | P15 Male KO vs P20 Male KO | **0.0448** |
| | P15 Male KO vs P30 Male KO | **n.s** |
| | P15 Female WT vs P15 Female KO | **n.s** |
| | P15 Female WT vs P20 Female WT | **n.s** |
| | P15 Female WT vs P30 Female WT | **n.s** |
| | P15 Female KO vs P20 Female KO | **0.0274** |
| | P15 Female KO vs P30 Female KO | **n.s** |
| | P20 Male WT vs P20 Male KO | **n.s** |
| | P20 Male WT vs P20 Female WT | **n.s** |
| | P20 Male WT vs P30 Male WT | **0.0337** |
| | P20 Male KO vs P20 Female KO | **n.s** |
| | P20 Male KO vs P30 Male KO | **0.0222** |
| | P20 Female WT vs P20 Female KO | **n.s** |
| | P20 Female WT vs P30 Female WT | **0.0135** |
| | P20 Female KO vs P30 Female KO | **0.0047** |
| | P30 Male WT vs P30 Male KO | **n.s** |
| | P30 Male WT vs P30 Female WT | **n.s** |
| | P30 Male KO vs P30 Female KO | **0.0416** |
| | P30 Female WT vs P30 Female KO | **n.s** |

**Table S10: Pair-wise post-hoc Fisher's group comparison following three-way RM ANOVA for response duration (PPI- startle alone trials).** Values in green indicate significant age difference and values in red indicate significant sex difference.

| Parameter | Group comparisons | Summary of significance |
|---|---|---|
| Response duration (PPI- startle alone trials) | P15 Male WT vs P15 Male KO | n.s |
| | P15 Male WT vs P15 Female WT | n.s |
| | P15 Male WT vs P20 Male WT | 0.0059 |
| | P15 Male WT vs P30 Male WT | 0.0249 |
| | P15 Male KO vs P15 Female KO | n.s |
| | P15 Male KO vs P20 Male KO | n.s |
| | P15 Male KO vs P30 Male KO | n.s |
| | P15 Female WT vs P15 Female KO | n.s |
| | P15 Female WT vs P20 Female WT | n.s |
| | P15 Female WT vs P30 Female WT | n.s |
| | P15 Female KO vs P20 Female KO | 0.0174 |
| | P15 Female KO vs P30 Female KO | n.s |
| | P20 Male WT vs P20 Male KO | n.s |
| | P20 Male WT vs P20 Female WT | n.s |
| | P20 Male WT vs P30 Male WT | n.s |
| | P20 Male KO vs P20 Female KO | n.s |
| | P20 Male KO vs P30 Male KO | 0.0386 |
| | P20 Female WT vs P20 Female KO | n.s |
| | P20 Female WT vs P30 Female WT | n.s |
| | P20 Female KO vs P30 Female KO | 0.0321 |
| | P30 Male WT vs P30 Male KO | n.s |
| | P30 Male WT vs P30 Female WT | 0.0379 |
| | P30 Male KO vs P30 Female KO | n.s |
| | P30 Female WT vs P30 Female KO | n.s |
